# ProteoPlotter: an executable proteomics visualization tool compatible with Perseus

**DOI:** 10.1101/2024.12.30.630796

**Authors:** Esther Olabisi-Adeniyi, Jason A. McAlister, Daniela Ferretti, Juergen Cox, Jennifer Geddes-McAlister

## Abstract

Mass spectrometry-based proteomics experiments produce complex datasets requiring robust statistical testing and effective visualization tools to ensure meaningful conclusions are drawn. The publicly-available proteomics data analysis platform, Perseus, is extensively used to perform such tasks, but opportunities to enhance visualization tools and promote accessibility of the data exist. In this study, we developed ProteoPlotter, a user-friendly, executable tool to complement Perseus for visualization of proteomics datasets. ProteoPlotter is built on the Shiny framework for R programming and enables illustration of multi-dimensional proteomics data. ProteoPlotter provides mapping of one-dimensional enrichment analyses, enhanced adaptability of volcano plots through incorporation of Gene Ontology terminology, visualization of 95% confidence intervals in principal component analysis plots using data ellipses, and customizable features. ProteoPlotter is designed for intuitive use by biological and computational researchers alike, providing descriptive instructions (i.e., Help Guide) for preparing and uploading Perseus output files. Herein, we demonstrate the application of ProteoPlotter towards microbial proteome remodeling under altered nutrient conditions and highlight the diversity of visualizations enabled with the platform for improved biological characterization. Through its comprehensive data visualization capabilities, linked to the power of Perseus data handling and statistical analyses, ProteoPlotter facilitates a deeper understanding of proteomics data to drive new biological discoveries.

## Introduction

Mass spectrometry (MS)-based proteomics allows for the comprehensive exploration of an organism’s proteome (i.e., composition of proteins within a biological system). Insights derived from powerful MS-based proteomics technology continue to bring advancements and innovation to diverse fields within the life and biomedical sciences ^1–4^. During proteomics experiments, MS instruments analyze protein mixtures across tens to thousands of samples through label-based and label-free approaches for detection, abundance, modifications, and interactions ^5^. To derive meaningful biological insights from the high-resolution data acquired, powerful computational platforms and search engines, such as MaxQuant, MSFragger, and DIA-NN ^6–8^, have been developed, applying comprehensive qualitative and quantitative analyses to identify and quantify peptides and proteins. The resulting data allows researchers to investigate the proteome, and apply statistical testing, to define and interpret important biological findings.

To facilitate the analysis of proteomics data and provide strategies for visualization and interpretation, software platforms for data visualization have been developed. These platforms range in utility and accessibility based on user expertise, experience, and expectations. For example, RStudio is a computational environment for the R programming language that offers several libraries for the statistical analysis of MS-based proteomics data but requires a working knowledge of R^9–11^. As a result, many researchers opt for computational programs which provide an intuitive graphical user interface (GUI) offering advanced features (e.g., statistical testing, normalization), with minimal requirement for computational expertise. This approach increases accessibility for biological researchers and facilitates robust approaches for proteomics data analyses. For example, Perseus, a powerful and versatile software that is freely available for the statistical analysis and visualization of proteomics datasets, is a widely-used and accessible platform ^12^. The Perseus GUI boasts an array of options for analyzing data through an interactive workflow that enables the user to track, preserve, and revisit a record of their activities on the platform. Some visualization options in Perseus include, but are not limited to, volcano plots, hierarchical clustering, and principal component analysis (PCA) plots.

To further harness the capabilities of the Perseus software, we introduce ProteoPlotter, an executable platform designed to utilize exported files from Perseus in generating multi-dimensional visualizations. Following user-enabled analysis in Perseus, such as filtering, normalization, computation of 1D annotation enrichment scores, and/or two-sample t-test, a raw tabular output can be exported from Perseus. ProteoPlotter utilizes these tabular datasets to transform the analyses into enhanced graphical representations. With its user-friendly GUI integrated within the Shiny web platform^13^, ProteoPlotter is designed to facilitate data visualization regardless of the user’s computational expertise to generate 1D annotation enrichment heat maps, interactive volcano plots, PCA plots with confidence intervales, dynamic range plots, Venn diagrams, and UpSet plots, within minimal effort. Here, we demonstrate application of ProteoPlotter to visualize proteome remodeling of the bacterial pathogen, *Klebsiella pneumoniae*, under altered iron conditions, providing new visualization of our biological findings ^14^. With ProteoPlotter, researchers can create multi-dimensional figures for MS data, effectively summarizing and conveying information for accessible and comprehensive data visualization and interpretation.

## Materials and Methods

### Software development and availability

ProteoPlotter is an interactive software designed to complement Perseus in the downstream analysis of quantitative proteomics data. ProteoPlotter was developed within the Shiny framework, based on the R programming language (version 4.3.2), incorporating several R libraries in its architecture, including ggplot2 (graphs), Shiny (GUI and web deployment), and Tidyverse (data structure design)^9,10,13,15,16^. The executable file to launch ProteoPlotter locally, with no RStudio or R software requirements, is available at https://github.com/JGM-Lab-UoG/. ProteoPlotter’s source code, which can be initiated within RStudio, is also available within the GitHub repository.

### ProteoPlotter Input

ProteoPlotter is specifically designed for all calculations to be performed within Perseus, taking full advantage of the Perseus data analysis workflow and archiving features. ProteoPlotter uses data contained within specific columns of the exported matrices to generate figures. The software accepts up to four Perseus matrices with specific plot-dependent column requirements, enabling users to generate the following data visualizations: (i) 1D annotation enrichment heat maps, (ii) multi-dimensional volcano plots, (iii) PCA plots with ellipses indicating 95% confidence intervals, (iv) dynamic range plots, (v) Venn diagrams of up to five groups, and (vi) UpSet plots (Table 1). All figures generated with ProteoPlotter are downloadable in high quality Portable Network Graphics (PNG) format. Additionally, the software features a ‘Guide’ tab with usage instructions. Helpful prompts also appear in the console as needed to assist users with troubleshooting.

**Table 1.** Perseus matrices and column requirements for upload to ProteoPlotter.

| Matrix uploaded to ProteoPlotter | Column headers in Perseus |  |  |  |  |  |
| --- | --- | --- | --- | --- | --- | --- |
|  | LFQ intensity <samples> | Majority protein IDs | Student t-test columns* | GO / Keywords columns<sup>+</sup> | x and y coordinates | 1D annotation columns<sup>~</sup> |
| 1D Annotation matrix |  |  |  |  |  | ✓ |
| Volcano, PCA and Dynamic Range matrix | ✓ | ✓ | ✓<br>(Optional: PCA and Dynamic Range plots) | Optional |  |  |
| Threshold Curve matrix |  |  |  |  | ✓ |  |
| Venn Diagram and UpSet matrix | ✓<br>(Mean intensity values) |  |  |  |  |  |

\*Student's t-test columns include: Significant, p-value, q-value, difference, and additional auto-generated columns.
<sup>+</sup>GO columns can include any or all of the following: Cellular Component, Molecular Function, Biological Process, and Keywords.
<sup>~</sup>1D annotation columns include the following: column, type, name, size, score, p-value, Benjamini-Hochberg False Discovery Rate <sup>17</sup>.

### 1D Gene Ontology Annotation Enrichment Heat Maps

Within Perseus, the user can perform a functional annotation of identified proteins, and subsequently, compute 1D annotation scores to investigate the enrichment of these annotation terms within samples^18^. Computed results are exported from Perseus as a matrix and uploaded to ProteoPlotter to generate 1-D enrichment heat maps. The matrix contains 1D annotation enrichment scores for included GO terms. Each 1D score is defined between −1 to 1 such that a value close to 1 correlates to the colour red on the heat map and indicates that the functional annotation term is enriched among proteins within one group of the comparative pair under consideration, while a value approaching −1 correlates to the colour blue, meaning that the annotation term is enriched among proteins within the other group of the comparative pair. A value near 0 is depicted as the colour white on the heat map and suggests non-preferential enrichment between the two groups. Within the ‘1D Annotation Heat Map’ tab in ProteoPlotter, the user uploads the Perseus matrix and up to four distinct heat maps are generated for (i) GO – cellular component (GOCC), (ii) GO – molecular function (GOMF), (iii) GO – biological process (GOBP), and (iv) UniProt Keywords. On each heat map, annotation terms are displayed along rows with columns indicating comparative pairs. Customizable options include: order of the groups, false discovery rate setting, score minimum/maximum values, and x-axis labels.

### Volcano plot

ProteoPlotter’s volcano plot feature allows the user to illustrate multiple layers of information from the dataset. For example, average protein intensity values and GO term annotations can be layered upon the plot (Figure 1). ProteoPlotter accepts up to two input matrices for the volcano plot, but the ‘Threshold Curve Matrix’ is optional; the ‘Matrix with t-test’ is required. The ‘Matrix with t-test’ input needs to contain the imputed protein intensities, Student’s t-test results, and majority protein identifications. When generating volcano plots for multiple comparative pairs, a separate t-test matrix should be generated for each pair. Subsequently, each matrix containing t-test results for a specific comparative pair is exported individually, rather than consolidating all t-test results into a single matrix. Furthermore, any of these t-test matrices can also serve as inputs for generating dynamic range or PCA plots given that the structure provides the required information regardless of the t-test performance.

**Figure 1:**
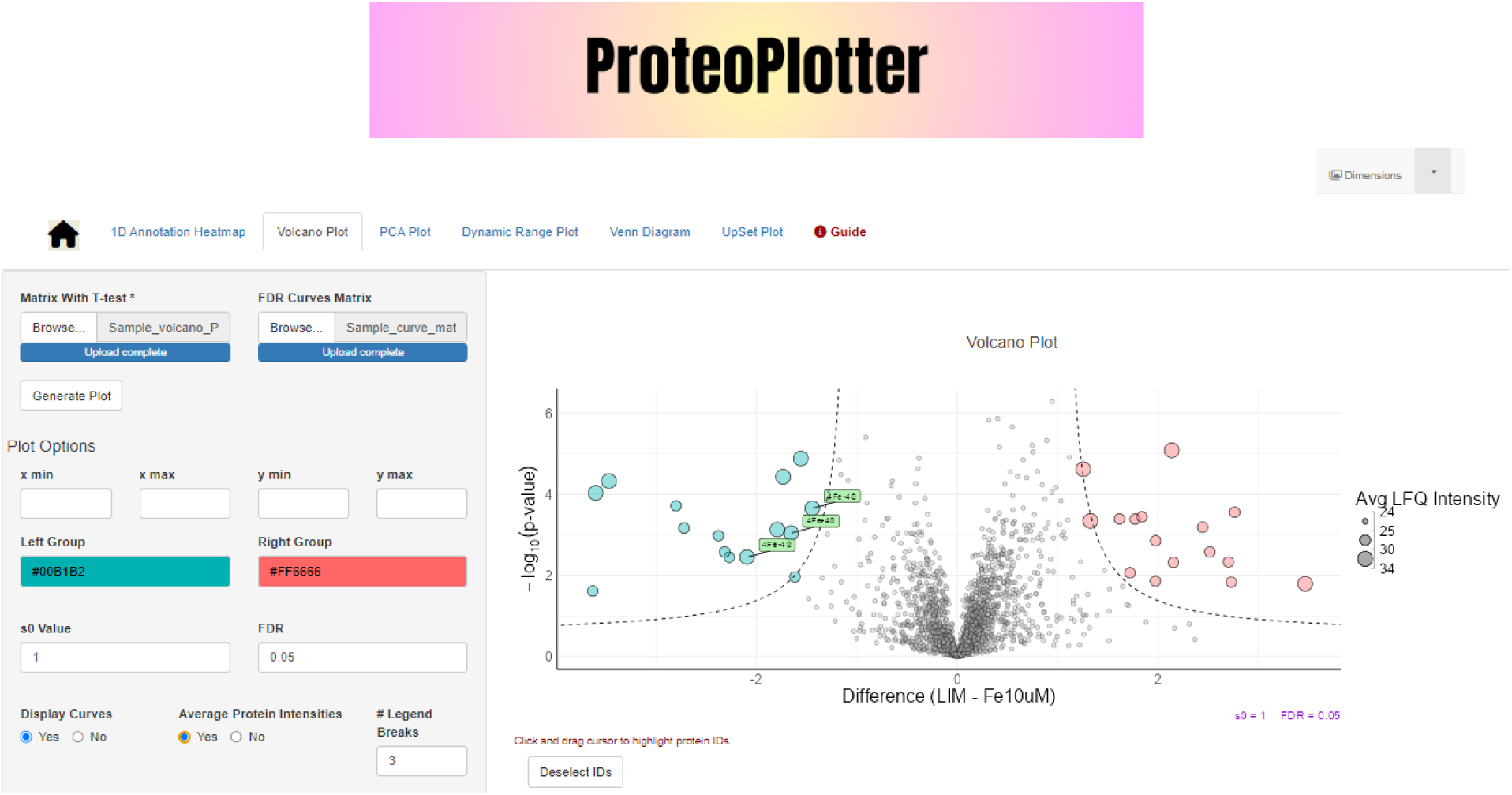
ProteoPlotter volcano plots. The size of each point (corresponding to a protein) varies by average protein abundance across replicates. The GO term annotation layer highlights the significantly different proteins annotated with a selected term.

On the volcano plot, negative log_10_ p-values and Student’s t-test differences from the matrix are plotted against each other. For depiction of functional annotations on the figure, the user must ensure that at least one functional annotation column (e.g., GOBP, GOCC, GOMF, Keywords) is provided. The user can also generate the False Discovery Rate (FDR) curves matrix in Perseus to layer FDR threshold curves on the volcano plot^17^. To achieve this, the user first creates a volcano plot in Perseus with the same S_0_ and FDR values that will be applied within ProteoPlotter. Subsequently, the FDR curves matrix containing the x and y coordinates is exported and uploaded. Additional volcano plot options within ProteoPlotter include hovering over points to disapply their respective GO terms within a chosen GO category, filtering GO terms to retain only those with 1D annotation scores (requires uploading the 1D annotation enrichment matrix), adjustment of axes limits, adjustment of FDR and S_0_ thresholds, and selection of colours from multiple colour palette libraries.

### Dynamic range and principal component analysis plots

As previously stated, the t-test matrix generated for the volcano plot can be used as an input file to generate dynamic range plots for all groups within the uploaded matrix, as well as a PCA plot. Alternatively, if volcano plots are not being generated, the user may export the matrix containing imputed protein intensities without performing t-tests from Perseus. Once the required matrix is uploaded, the ‘Generate Plot’ button within the ‘Dynamic Range Plot’ tab or the ‘PCA Plot’ tab can be used to generate the figures. The dynamic range plots illustrate the average protein abundance (log_10_) across replicates against their descending-order ranking, depicting the distribution of protein abundance across measured proteomes. With ProteoPlotter’s traditional PCA function, users can investigate clustering patterns among group replicates and the variance driving separation among groups. Moreover, the software also offers the option to overlay data ellipses onto the PCA plot to illustrate the multivariate t-distribution of the data at a 95% confidence interval^19,20^. Note that this data ellipse feature is only applicable to groups with four or more replicates.

### Venn diagram and UpSet plots

ProteoPlotter generates Venn diagrams and UpSet plots from average protein intensity values computed in Perseus. To produce a Venn diagram, the user uploads the matrix within the ‘Venn Diagram’ tab, then selects the ‘Generate Plot’ button. The Venn diagram allows users to assess unique and shared proteins across up to five groups, and users can select colours for each group within the Venn diagram. In contrast, the UpSet plot in ProteoPlotter largely overcomes this limitation of group number by enabling users to visualize up to 25 groups using the same matrix uploaded for the Venn diagram. With the matrix already uploaded, the user selects the ‘Generate Plot’ button within the ‘UpSet Plot’ tab to produce the figure.

### Analysis of Sample Dataset

A dataset from a previous proteomics study on the bacterial pathogen, *Klebsiella pneumoniae*, was selected to demonstrate the multi-dimensional visualization features available within ProteoPlotter^14^. The dataset was obtained from the PRIDE database (Project PXD015623)^21^. In the study, the impact of iron limitation on bacterial growth, survival, and proteome differentiation of *K. pneumoniae* was analyzed. The bacterium was cultured in low iron media, and media supplemented with Fe_2_(SO_4_)_3_ at 10 µM and 100 µM in four biological replicates for each condition. Cellular proteome samples were obtained from each treated group using tandem mass spectrometry techniques (LC-MS/MS) as previously described^14^. Raw mass spectrometry files were processed with MaxQuant (version 1.6.0.26), and the tab-delimited output from MaxQuant, proteingroups.txt, was imported into Perseus (version 2.1.3.0) (Figure 2).

**Figure 2:**
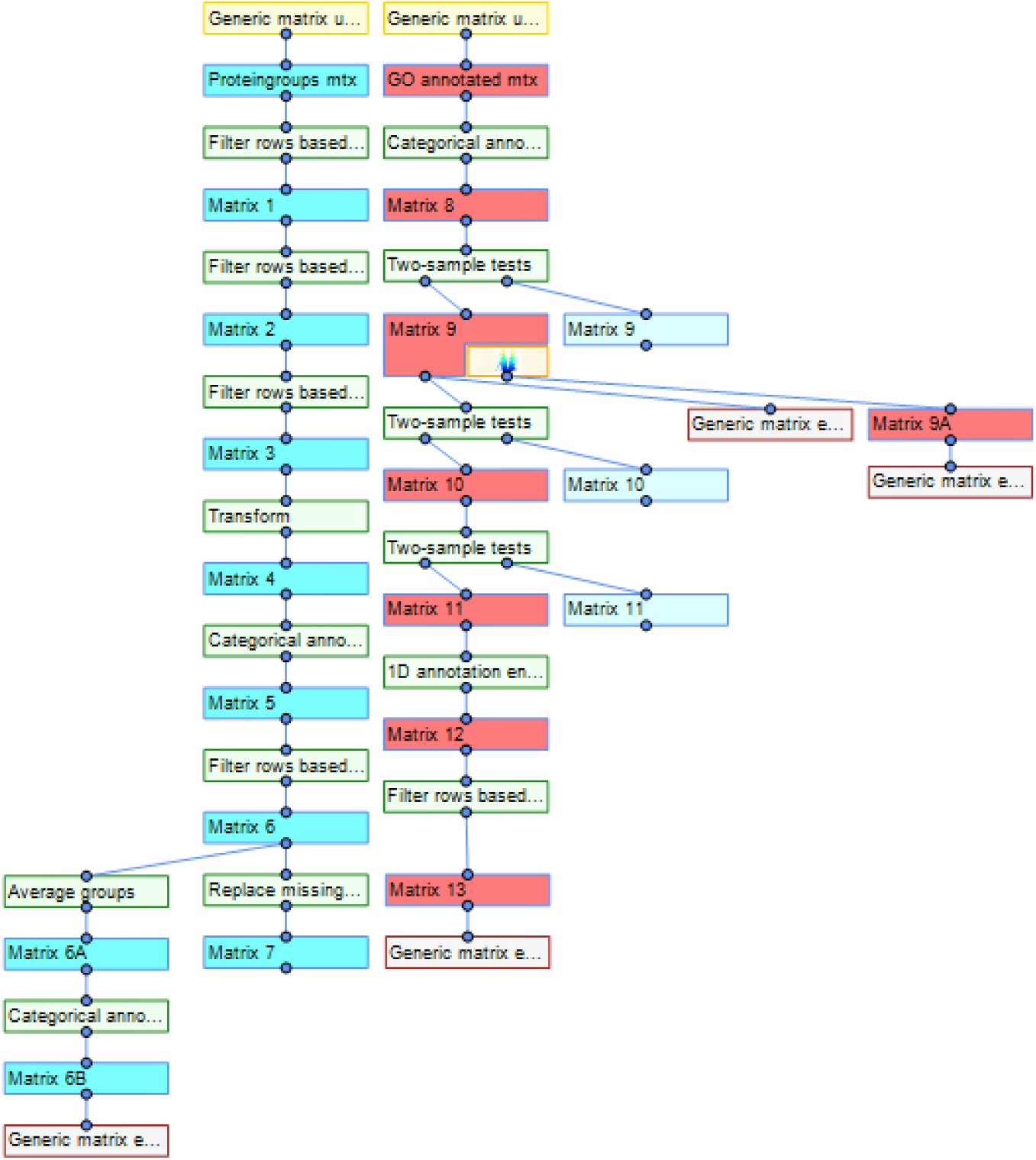
Perseus workflow for cellular proteome analysis of *K. pneumoniae* under altered iron conditions. Teal workflow: before GO annotation; Red workflow: after GO annotation.

The workflow in Perseus performed the following analyses, including removal of proteins that matched to the reverse database, contaminants, and proteins only identified with modified peptides (matrices 1-3). Intensities were converted to a log_2_ scale (matrix 4), and LFQ intensity columns were categorically annotated as follows: LIM, Fe10 µM, and Fe100 µM (matrix 5). Only proteins with a minimum of three valid values within at least one group were retained to filter the total number of proteins identified (matrix 6). Next, we averaged intensity values to derive the matrix for the Venn diagram and UpSet plot, applying a minimum of three valid values per group as the threshold (matrix 6A). The resulting matrix was reannotated with group names and exported (matrix 6B). Subsequently, missing values were imputed with numbers drawn from a normal distribution with a downshift by 1.8 standard deviations and a width of 0.3 standard deviations (matrix 7). GO and UniProt Keywords annotations were collected from UniProtKB using the UniProt ID mapper feature^22,23^. The annotations were added to the matrix outside of the Perseus platform and the annotated matrix was re-uploaded to the Perseus workflow (matrix ‘GO annotated).

For statistical testing, Student’s t-test (two-sample test) were computed to identify proteins with significant differences in abundance (p-value ≤ 0.05) between LIM and Fe10 µM; LIM and FE100 µM; Fe10 µM and Fe100 µM (matrix 9-11). Permutation-based FDR was applied for multiple hypothesis testing correction with 250 randomizations, FDR = 0.05, and S_0_ = 1. To use the ProteoPlotter volcano plot feature, we exported the matrix containing the Student’s t-test results for LIM vs. Fe10 µM (matrix 9). We used the same matrix to create both the PCA and dynamic range plots. Additionally, threshold curves for the volcano plots were generated by exporting the curve matrix produced by the volcano plot feature in Perseus (matrix 9A). For the 1D annotation heat maps, we computed 1D annotation scores suing the ‘t-test difference’ columns of all three comparative pairs present within matrix 11 (FDR = 0.05). We also filtered the resulting matrix (matrix 12) to exclude rows with non-annotation terms. The final matrix was exported to ProteoPlotter to generate the 1D annotation heat maps (matrix 13). All matrices were exported in the .txt format.

## Results

### Quantitative characterization of the K. pneumoniae cellular proteome under altered iron conditions using ProteoPlotter

We applied ProteoPlotter’s volcano plot feature for exploring the proteins that were significantly differentially abundant between LIM and FE10 µM conditions (Figure 3A). A significant difference was observed for 31 proteins; 16 proteins were significantly more abundant under iron-limited (LIM) conditions while 15 proteins were significantly more abundant under iron-replete (10 µM Fe_2_(SO_4_)_3_) conditions. The S_0_ and FDR values, which are adjustable when creating the plot, are displayed in purple at the base of the plot for the user’s information. Users can choose to include or exclude the threshold curves.

**Figure 3:**
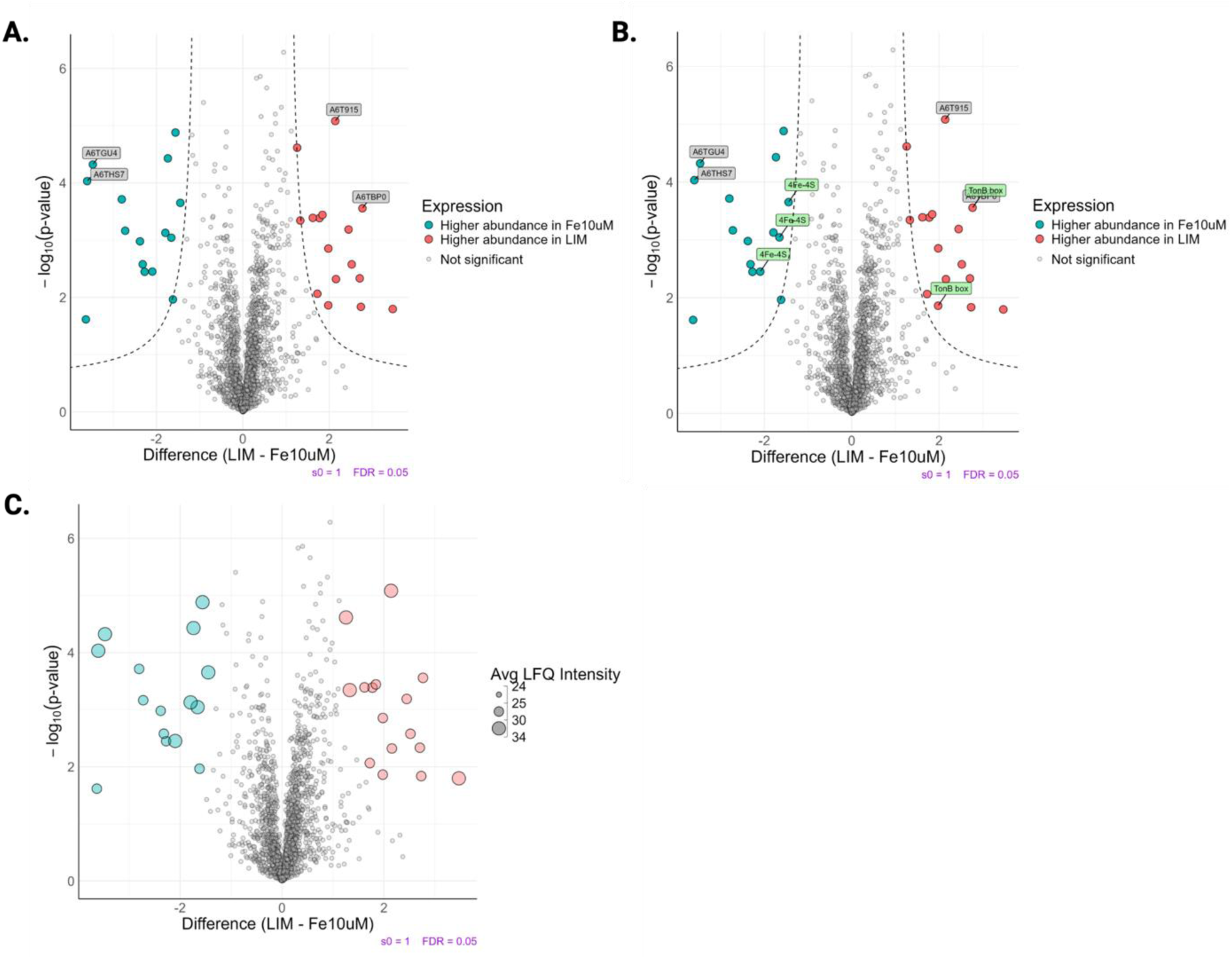
Significantly different protein abundance profiles under iron-limited (LIM) vs. iron-replete (Fe10 µM) conditions using ProteoPlotter’s volcano plot feature. A) Protein ID selection layer labeling select proteins. B) GO annotation layer displaying the significantly different proteins associated with TonB box, a GO term under the Keywords category. C) Volcano plot layer with average intensity values for significantly different proteins in each treatment group. Student’s t-test p-value < 0.05; FDR = 0.05; S_0_ = 1.

ProteoPlotter also allows the user to highlight proteins IDs, enabling rapid identification of specific significantly different proteins. Next, we overlaid select UniProt Keywords onto the already labeled volcano plot, thereby allowing us to visualize and investigate the functional properties of significantly different proteins (Figure 3B). For example, we illustrate the feature by labeling GO annotations associated with the TonB box, a protein involved in ferric-siderophore receptor function to provide a mechanical linkage between the cytoplasmic and outer membrane, promoting energy transduction from the proton motive force to the receptor^24–27^. Other GO categories can also be illustrated in ProteoPlotter. In addition, we visualized the average protein intensity for each significantly altered protein. Through this figure, we observed that most of the significantly differentially abundant proteins in the iron-limited samples exhibited relatively moderate intensity levels (Figure 3C). In contrast, the Fe10 µM samples demonstrated a consistent distribution of both low and high intensity values.

Using ProteoPlotter, we generated a 1D annotation heat map based on UniProt Keywords to gain another perspective into the functional enrichment of the cellular proteome under all three tested conditions (Figure 4). Several categories showed positive enrichment in replete (Fe10 µM and Fe100 µM) conditions compared to limited conditions, including rRNA-binding, ribosomal, and ribonucleoprotein categories. We also observed a positive enrichment of iron transport and TonB GO terms under limited conditions relative to Fe100 µM replete conditions. More iron-associated terms, such as iron-sulfur and 4Fe-4S clusters were positively enriched under 10 µM replete conditions relative to 100 µM replete conditions. The patterns observed in the heat maps suggest that changes in iron availability alters the cellular environment in *K. pneumoniae*, leading to an increase in iron-associated protein production under low iron conditions.

**Figure 4:**
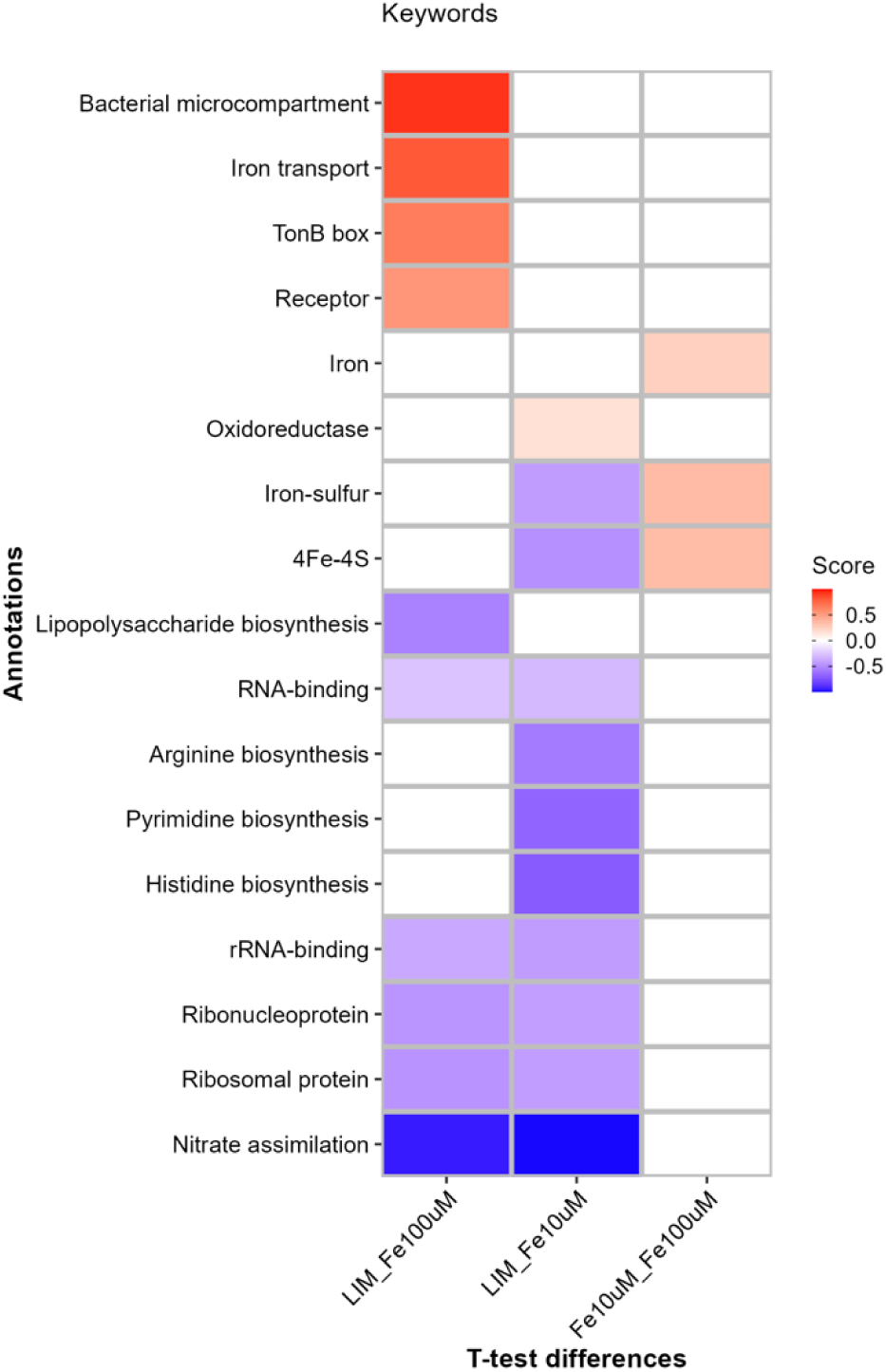
1D annotation enrichment heat map for UniProt Keywords for *K. pneumoniae* under altered iron conditions. Student’s t-test p-value < 0.05; FDR = 0.05; −1.0 < Score < 1.0.

### Qualitative characterization of the K. pneumoniae cellular proteome under altered iron conditions using ProteoPlotter

The Venn diagram in ProteoPlotter illustrates the shared and unique number of proteins identified among groups, as counts, percentages or both (Figure 5A). The majority of identified proteins are defined within a core proteome of 1855 proteins present across all three tested conditions. Notably, 33 proteins were unique to Fe100 µM and 36 proteins were unique to Fe10 µM conditions, whereas the iron-limited conditions demonstrated unique identification of 52 proteins. This information is also depicted using the UpSet plot feature (Figure 5B).

**Figure 5:**
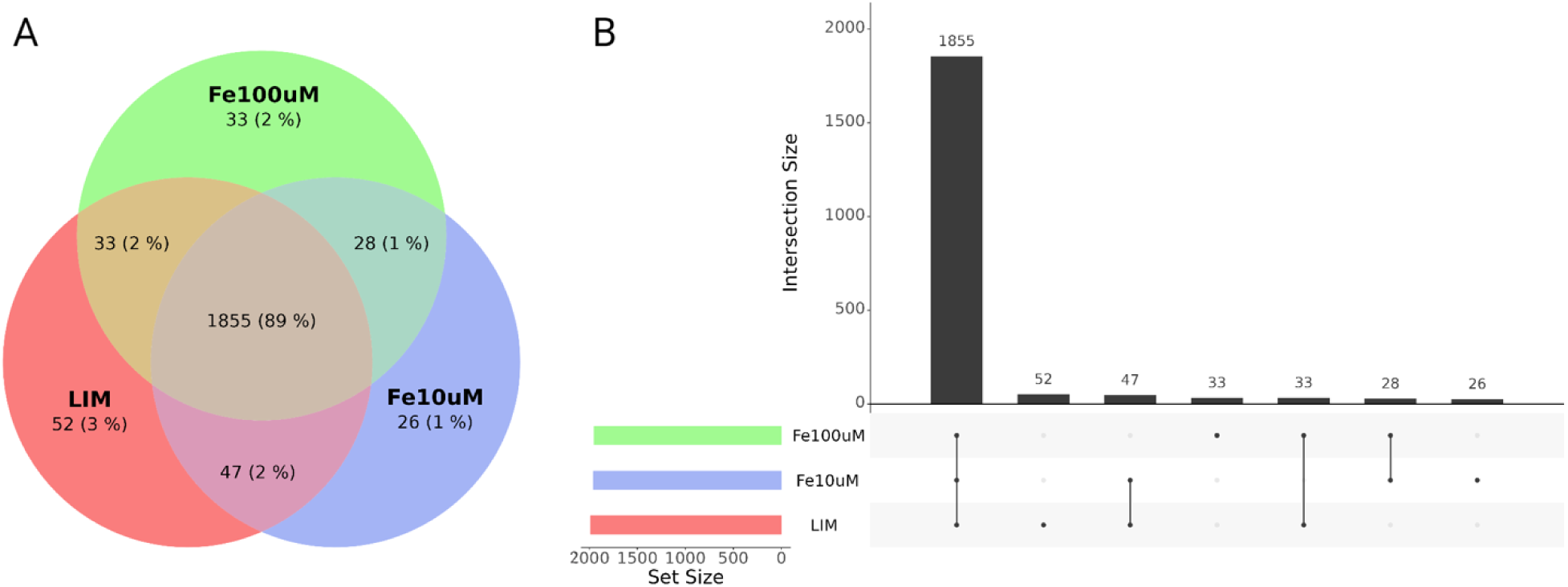
Qualitative analysis Venn and UpSet plots in ProteoPlotter. A) Venn diagram comparing the number of proteins identified in the cellular proteome cultured under limited (LIM) and replete (Fe10 µM and Fe100 µM) iron conditions. B) UpSet plot demonstrating protein identification patterns across the conditions.

Next, we generated PCA and dynamic range plots using imputed LFQ intensities, to explore the distribution of protein abundance values. In the PCA, treatment groups clustered, with the largest variability observed between the iron-limited group and both iron-replete groups (Principal Component 1 = 22.78%; Figure 6A). The second principal component accounted for 16.64% of the variance, reflecting the reproducibility among replicates. We also added data ellipses to the PCA plot to explore the t-distribution of samples within each treatment group. This step revealed an overlap between both replete conditions and an isolated ellipse for the iron-limited samples (95% CI; Figure 6B). Lastly, dynamic range plots were generated to assess the varying abundance of proteins identified in the cellular proteome under iron-limited and iron-replete conditions. The protein abundance within each group exhibited a dynamic range of distribution, with proteins showing high, moderate, and low intensities, all spanning across a similar range of 8 to 11 (log_10_ LFQ intensity) (Figure 7).

**Figure 6:**
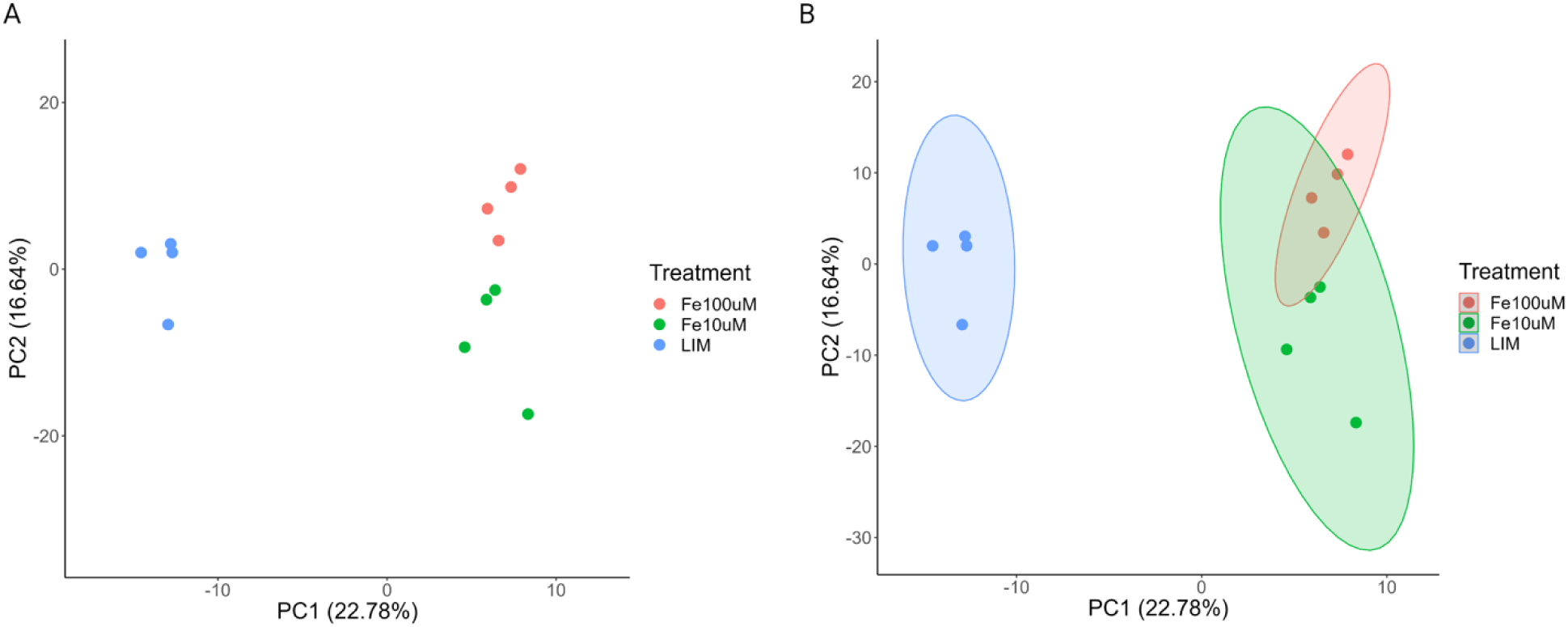
Qualitative analysis PCA plots in ProteoPlotter. A) PCA of the cellular proteome identifying clusters by group. B) Data ellipses visualizing the multivariate t-distribution of each treatment group (95% CI).

**Figure 7:**
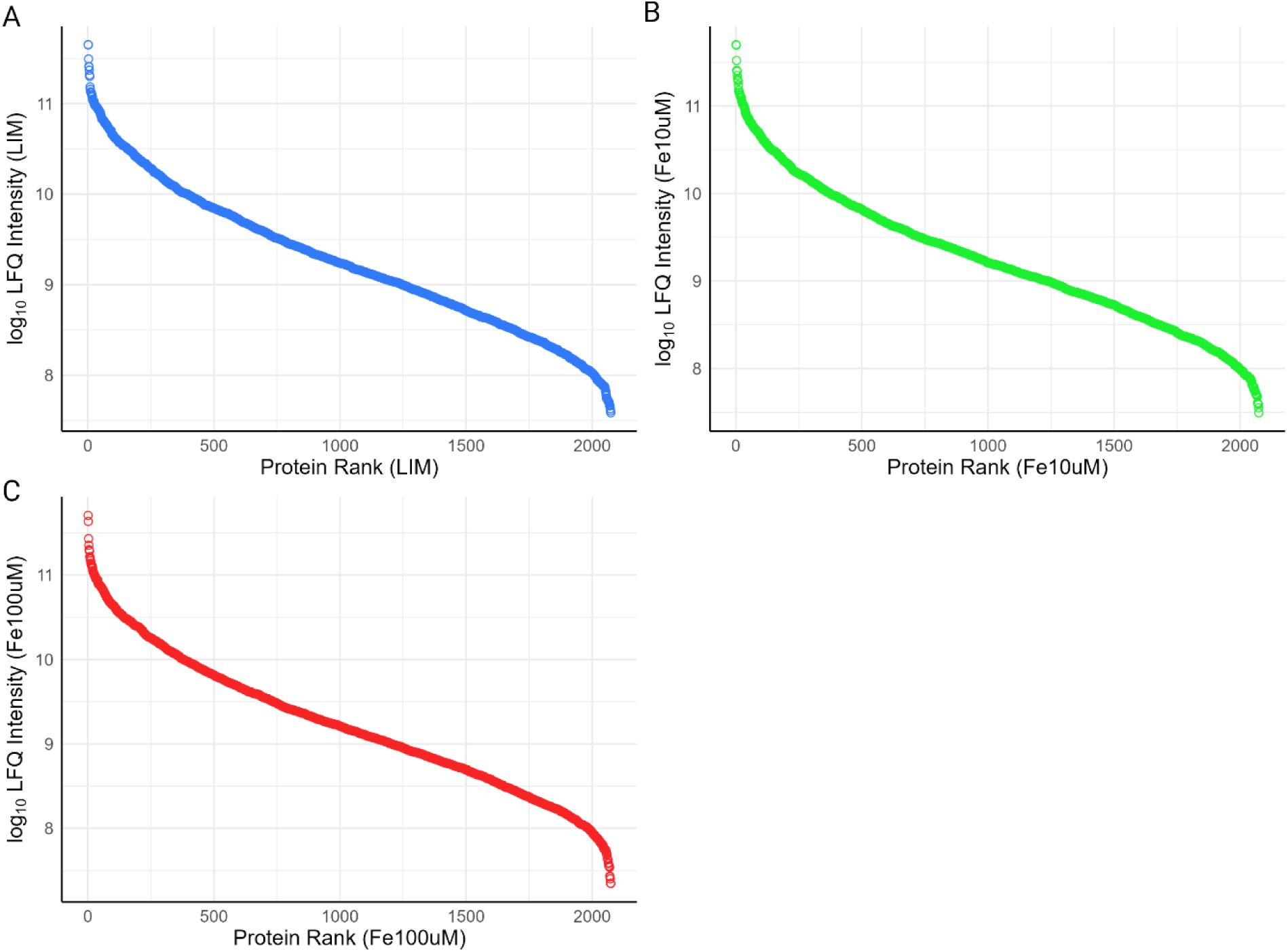
Qualitative analysis dynamic range plots in ProteoPlotter. Average protein abundance plotted against protein ranks (highest to lowest) to illustrate the dynamic range of *K. pneumoniae* cellular proteome under iron-limited and iron-replete conditions.

## Discussion

ProteoPlotter provides new visualization tools and capabilities, and application to a previously published proteomics dataset of *K. pneumoniae* results in a demonstrated ability to decipher insights into how iron availability affects the bacteria’s cellular proteome. We selected proteins highlighted in the volcano plot and identified them with the UniProtKB database to illustrate our ability to use ProteoPlotter for investigation of biologically relevant proteins. For instance, CirA (A6TBP0), an outer membrane pore protein requiring TonB, was present at significantly higher abundance under iron-limited conditions ^14,27^. CirA associated with siderophore secretion, which is demonstrated by *K. pneumoniae* to improve iron uptake under low-iron availability^28,29^. Gram-negative bacteria, such as *Klebsiella* spp., use siderophore-release mechanisms under iron-limited conditions as the molecule’s high affinity for metals improves iron uptake^30,31^. Therefore, an increase in CirA’s abundance likely reflects the activation of siderophore-release mechanisms triggered by low-iron conditions. Another protein, Fe(II)-2OG-dependent oxygenase (A6T915) was also more abundant under iron-limited conditions, which is consistent with previous findings that show a correlation between increased Fe(II)-2OG-dependent oxygenase activity in *Saccharomyces cerevisiae* and low iron conditions^32^.

ProteoPlotter’s multidimensional approach to data visualization enhances data analysis and interpretation of results. For instance, overlaying GO terms onto the volcano plot revealed that CirA protein is associated with the TonB box annotation term. We also observed a positive enrichment of TonB under iron-limited conditions in the 1D heat map, which is likely correlated with the significantly higher abundance of CirA, a protein that is dependent on TonB. Analyzing this dataset from multiple perspectives through the volcano plot and heat map enables the user to corroborate associations between a protein and defined biological functions. To further investigate GO terms associated with significantly different proteins, ProteoPlotter users can focus on key terms by applying the filter that retains only those terms enriched in the 1D annotation dataset.

During proteomics analysis, it is crucial for proteins to be detectable across varying abundances, particularly for low abundant proteins^33^. Through the dynamic range plot in ProteoPlotter, we observed a well distributed spectrum of protein abundances, including those at lower levels, across five orders of magnitude. Researchers can apply this tool before proceeding with in-depth analysis and visualization. Additional analysis, with the Venn diagram and UpSet plot features indicates that the cellular proteome under iron-limited conditions contained more unique proteins relative to the proteome within iron-replete environments. Given the significantly higher abundance of siderophore-associated proteins like CirA under iron-limited conditions, it is likely that iron-limited samples produce more condition-specific proteins to facilitate efficient siderophore-release and iron uptake. Dimensionality reduction with PCA in ProteoPlotter also revealed a clear distinction between the replicates in iron-limited environments and those in iron-replete environments. The clusters for both 10 µM and 100 µM samples were positioned closely, suggesting a similarity between the proteomic profiles and a separation from the iron-limited cluster. T-distribution ellipses of the groups further demonstrate that the iron-limited proteome of distinct from both iron-replete proteomes. From the overlapping ellipses, we can infer that both iron-replete samples maintain similarity to one another whereas the proteomic profile under limited conditions remains distinct (95% CI).

## Conclusion

ProteoPlotter serves as a valuable supplementary visualization tool for proteomics data analyzed in Perseus to offer new modes of extracting biological insights from complex data. The program enabled us to successfully illustrate, and extract details embedded within our sample data. ProteoPlotter integrates multiple data visualization techniques into a user-friendly, executable platform, allowing for a harmonized exploration of data in tandem with Perseus. Presenting results graphically and exploring traditional visualization methods in a multidimensional manner empowers researchers to gain a clearer understanding of their data, thereby improving data interpretation.

## Data availability statement

The executable file to launch ProteoPlotter locally, with no RStudio or R software requirements, is available at https://github.com/JGM-Lab-UoG/. ProteoPlotter’s source code, which can be initiated within RStudio, is also available within the GitHub repository. Proteomics data is available through PRIDE database (Project PXD015623).

## Acknowledgements

The authors thank members of the Geddes-McAlister lab for testing of ProteoPlotter and insightful discussions and feedback throughout the development process.

## Contributions

J.A.M. & J.G.-M. conceptualized the design and development of ProteoPlotter. E.O.-A. designed and developed ProteoPlotter. D.F. & J.C. provided guidance and input into the design and development of ProteoPlotter. E.O.-A. & J.A.M. prepared the first draft of the manuscript. E.O.-A. & J.G.-M. prepared figures. J.A.M. & J.G.-M. prepared the final version of the manuscript. All authors have reviewed and approve the submitted manuscript.

